# VAE-Sim: a novel molecular similarity measure based on a variational autoencoder

**DOI:** 10.1101/2020.06.26.172908

**Authors:** Soumitra Samanta, Steve O’Hagan, Neil Swainston, Timothy J. Roberts, Douglas B. Kell

**Author notes:** These authors contributed equally (Soumitra Samanta, Steve O’Hagan). University College London Hospital NHS Foundation Trust, 250 Euston Road, London, NW1 2PB.

## Abstract

Molecular similarity is an elusive but core ‘unsupervised’ cheminformatics concept, yet different ‘fingerprint’ encodings of molecular structures return very different similarity values even when using the same similarity metric. Each encoding may be of value when applied to other problems with objective or target functions, implying that *a priori* none is ‘better’ than the others, nor than encoding-free metrics such as maximum common substructure (MCSS). We here introduce a novel approach to molecular similarity, in the form of a variational autoencoder (VAE). This learns the joint distribution p(z|x) where z is a latent vector and x are the (same) input/output data. It takes the form of a ‘bowtie’-shaped artificial neural network. In the middle is a ‘bottleneck layer’ or latent vector in which inputs are transformed into, and represented as, a vector of numbers (encoding), with a reverse process (decoding) seeking to return the SMILES string that was the input. We train a VAE on over 6 million druglike molecules and natural products (including over one million in the final holdout set). The VAE vector distances provide a rapid and novel metric for molecular similarity that is both easily and rapidly calculated. We describe the method and its application to a typical similarity problem in cheminformatics.

## Introduction

The concept of molecular similarity lies at the core of cheminformatics [1-3]. It implies that molecules of ‘similar’ structure tend to have similar properties. Thus, a typical question can be formulated as follows: “given a molecule of interest M, possibly showing some kind of chemical activity, find me the nearest 50 molecules from a potentially huge online collection to purchase that are most similar to M so I can assess their behaviour in a relevant quantitative-structure-activity (QSAR) analysis”.

The most common strategies for assessing molecular similarity involve encoding the molecule as a vector of numbers, such that the vectors encoding two molecules may be compared according to their Euclidean or other distance. In the case of binary strings the Jaccard or Tanimoto similarity (TS) is commonly used [4] as it is a metric (between zero and one). One means for obtaining such a vector for a molecule is to calculate from the structure (or measure) various <u>properties</u> of the molecule (‘descriptors’ [5-7]), such as clogP or total polar surface area, and then to concatenate them. However, a more common strategy for obtaining the encoding vector of numbers is simply to use structural features <u>directly</u> and to encode them as so-called molecular fingerprints [8-17]. Well-known examples include MACCS [18], atom pairs [19], torsion [20], extended connectivity [21], functional class [22], circular [23], and so on. The similarities so encoded can also then be compared as their Jaccard or Tanimoto similarities. Sometimes a ‘difference’ or ‘distance’ is discussed and formulated as 1-TS (a true metric). An excellent and widely used framework for doing all of this is RDKit (www.rdkit.org/) [24], that presently contains nine methods for producing molecular fingerprints.

The problem comes from the fact that the ‘most similar’ molecules to a target molecule often differ wildly both as judged by their structures observable by eye and quantitatively in terms of the value of the TS of the different fingerprints [25]. As a very small and simple dataset, we take the set of molecules observed by Dickens and colleagues [26] to inhibit the transporter-mediated uptake of the second-generation atypical antipsychotic drug clozapine. These are Olanzapine, Chlorpromazine, Quetiapine, Prazosin, Lamotrigine, Indatraline, Verapamil and Rhein. Of the FDA approved drugs, we assessed the top 50 drugs in terms of their structural similarity to clozapine using the nine RDKit encodings, with the results shown in Table 1 and Fig 1. Only the first four of these are even within the top 50 for any encoding, and only olanzapine appears for each of them. By contrast, the most potent inhibitor is prazosin (which is not even wholly of the same drug class, being both a treatment for anxiety and a high-blood-pressure-lowering agent); however, it appears in the top 50 in only one encoding (torsion) and then with a Tanimoto similarity of just 0.37. This said, visual inspection of their ‘Kekularised’ structures does show a substantial common substructure between prazosin and clozapine (marked in Fig 1). It is clear that the similarities as judged by standard fingerprint encodings are <u>highly variable</u>, and are prone to both false negatives and false positives when it comes to attacking the question as set down above. What we need is a different kind of strategy.

**Table 1.** Tanimoto similarity to clozapine using nine different RDKit encodings and their ability to inhibit clozapine transport (data extracted from [26]). A shaded cell means that the molecule was not judged to be in the ‘top 50’ using that encoding.

| Drug | % inhib<br>clozapine<br>uptake | TS Atom<br>Pair | TS<br>Avalon | TS Feat<br>Morgan | TS<br>layered | TS<br>MACCS | TS<br>Morgan | TS<br>Pattern | TS<br>RDKit | TS<br>Torsion |
| --- | --- | --- | --- | --- | --- | --- | --- | --- | --- | --- |
| Olanzapine | 41 | 0.68 | 0.47 | 0.55 | 0.77 | 0.8 | 0.53 | 0.81 | 0.74 | 0.66 |
| Chlorpromazine | 75 | 0.53 |  | 0.35 |  | 0.66 | 0.3 | 0.74 |  | 0.33 |
| Quetiapine | 65 | 0.51 | 0.57 | 0.42 | 0.78 |  | 0.35 | 0.8 |  | 0.48 |
| Prazosin | 94 |  |  |  |  |  |  |  |  | 0.37 |
| Lamotrigine | 26 |  |  |  |  |  |  |  |  |  |
| Indatraline | 35 |  |  |  |  |  |  |  |  |  |
| Verapamil | 83 |  |  |  |  |  |  |  |  |  |
| Rhein | 39 |  |  |  |  |  |  |  |  |  |

**Figure 1.**
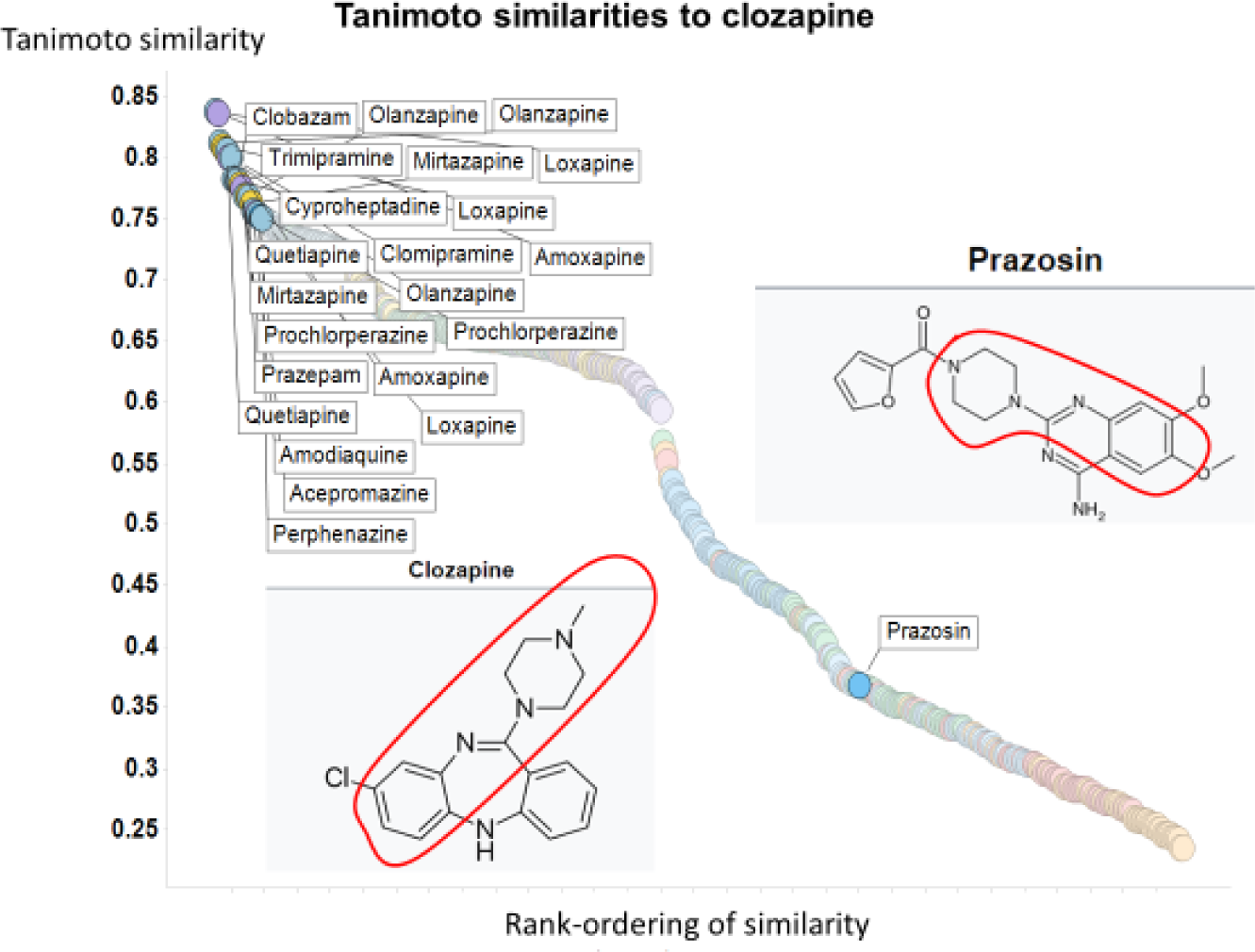
Tanimoto similarities of various molecules to clozapine using the Torsion encoding from RDKit.

The typical structure of a QSAR type of problem is given in Fig 2A, where a series of molecules represented as SMILES strings [27] are encoded as molecular fingerprints and used to learn a nonlinear mapping to produce an output in the form of a classification or regression estimation. The architecture of this is implicitly in the form of a multilayer perceptron (a classical neural network [28-31]), in which weights are modified (‘trained’) to provide a mapping from input SMILES to a numerical output. Our fundamental problem stems from the fact that these types of encoding are one-way: The SMILES string can generate the molecular fingerprint but the molecular fingerprint cannot generate the SMILES. Put another way, it is the transition from a world of discrete objects (here molecules) with categorical representations (here SMILES strings) to one of a continuous representation (vectors of weights) that is seemingly irreversible in this representation. One key element is the means by which we can go from a non-numerical representation (such as SMILES or similar [32]) to a numerical representation or ‘embedding’ (that, as we shall see, is typically constituted by vectors of numbers in the nodes and weights of multilayer neural networks) [33-38]). Deep learning has also been used for the <u>encoding</u> step of 2D chemical structures [39; 40].

**Figure 2.**
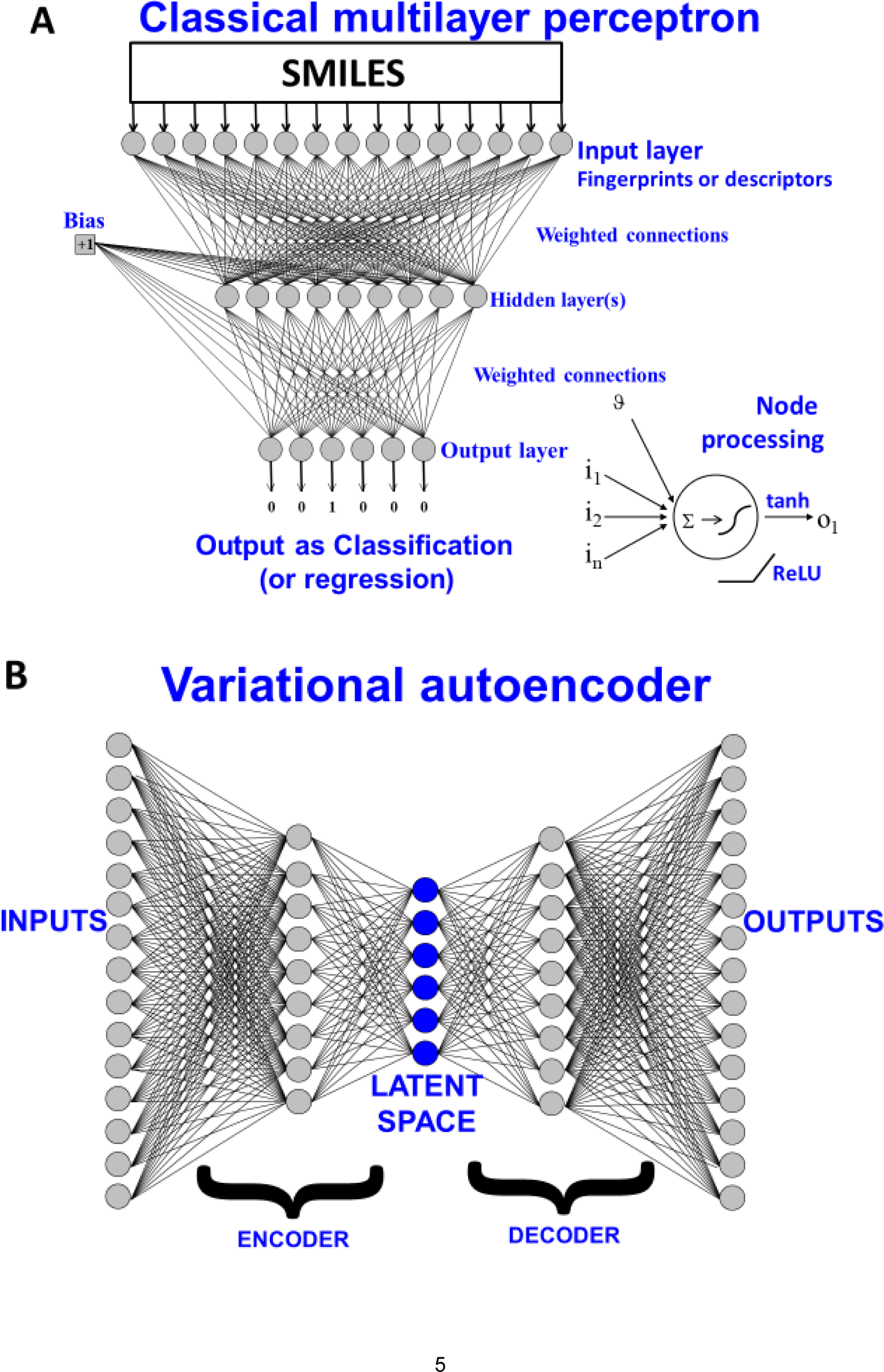
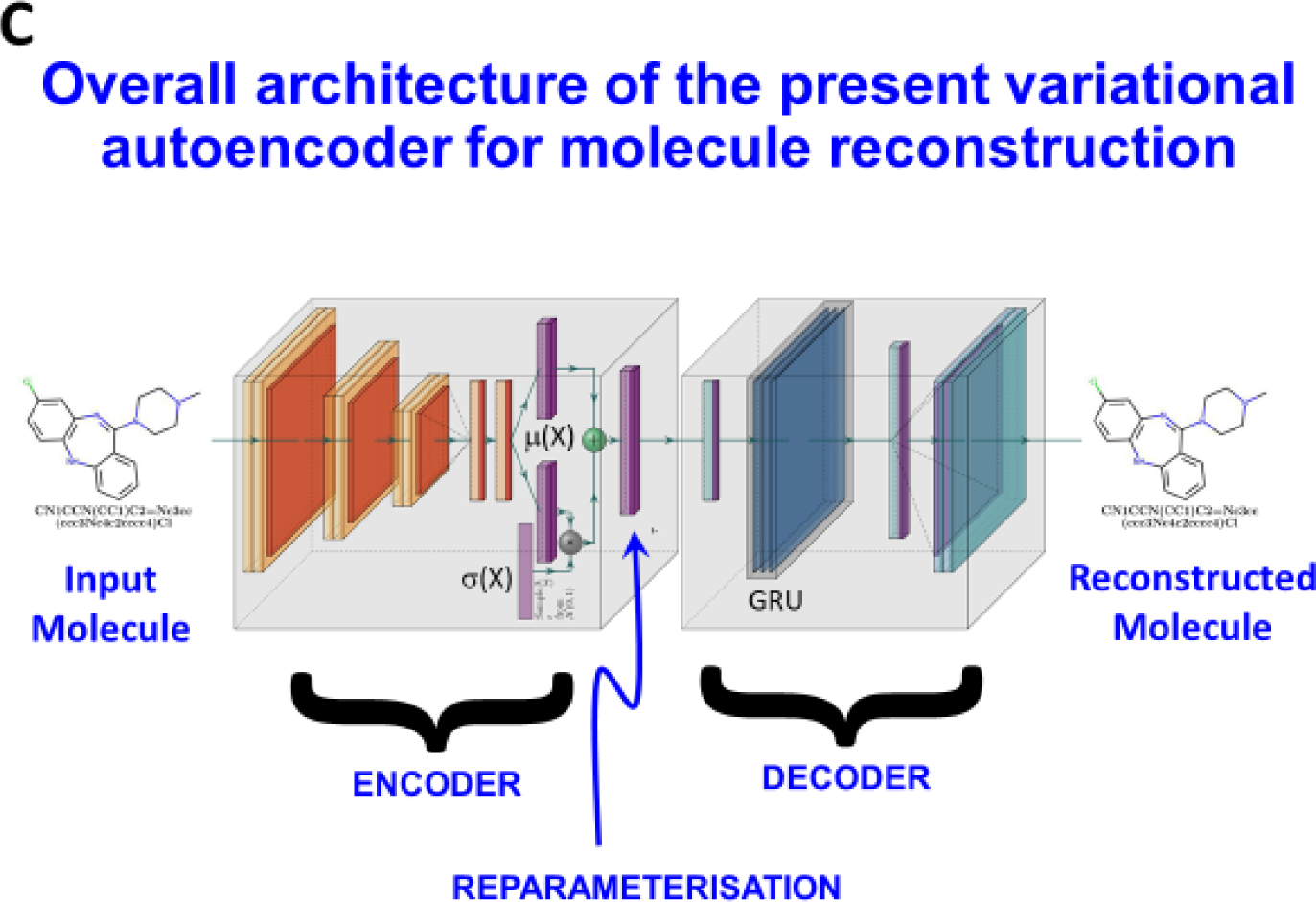
Two kinds of neural architecture. **A**. A classical multilayer perceptron representing a supervised learning system in which molecules encoded as SMILES strings can be used as paired inputs with outputs of interest (whether a classification or a regression). The trained model may then be interrogated with further molecules and the output ascertained. **B**. A variational autoencoder, is a supervised means of fitting distributions of discrete models in a way that reconstructs them via a vector in a latent space. **C**. The VAE architecture used in the present work.

More recently, it was recognised that various kinds of architectures could in fact permit the reversal of this numerical encoding so as to return a molecule (or its SMILES string encoding a unique structure). These are known as <u>generative methods</u> [41-50], and at heart their aim to generate a suitable and computationally useful <u>representation</u> [51] of the input data. It is common (but cf. [52; 53]) to contrast two main flavours: generative adversarial networks [54-61] and (especially variational) autoencoders (VAEs) [41; 42; 62-71]. We focus here on the latter, illustrated in Fig 2B. VAEs are latent-variable generative models that define a joint density p_θ_(x,z) between some observed data 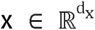 and unobserved or latent variables 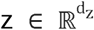 [72], given some model parameters θ. They use a variational posterior (also referred to as an encoder), q_φ_(z | x), to construct the latent variables with variational parameters φ, and a combination of p(z) and p(x|z) to create a decoder that has the opposite effect. Learning the posterior directly is computationally intractable, so the generic deep learning strategy is to train a neural network to approximate it. The original ‘error’ backpropagated was based on the Kullback-Leibler (KL) divergence between the desired (log likelihood reconstruction error) and the predicted output distributions [62]. A very great many variants of both architectures and divergence metrics have been proposed since then (not all discernibly better [73]), and it is a very active field (e.g. [58; 59; 74; 75]). Since tuning is necessarily domain-specific [76], and most work is in the processing of images and natural languages rather than in molecules, we merely mention a couple, such as transformers (e.g. [77; 78]) and others (e.g. [79; 80]). Crucial to such autoencoders (that can also be used for data visualisation [81]) is the concept of a bottleneck layer, that as a series of nodes of lower dimensionality than its predecessors or successors, serves to extract or represent [51] the crucial features of the input molecules that are nonetheless sufficient to admit their reconstruction. Indeed, such strategies are sometimes referred to as representational learning.

A higher-level version of the above might state that a good variational autoencoder will project a set of discrete molecules into a continuous latent space represented for any given molecule by the vector representing the values of the outputs of the nodes in the bottleneck layer when it (or its SMILES representation) is applied to the encoder as an input. As with the commonest neural net training system (but cf. [82-86]), we use backpropagation to update the network so as to minimise the difference between the predicted and the desired output, subject to any other constraints that we may apply. We also recognise the importance of various forms of regularisation, that are all designed to prevent overfitting [49; 87-90].

Because the outputs of the nodes in the bottleneck layer both (i) encode the molecule of interest and (ii) effectively represent where molecules are in the chemical space on which they have been trained, a simple metric of similarity between two molecules is clearly the Euclidean or other comparable distance (e.g. cosine distance) between these vectors. This thus provides for a novel type of similarity encoding, that in a sense relates the whole chemical space on which the system has been trained and that we suspect may be of general utility. We might refer to this encoding as the ‘essence of molecules’ (EM) encoding, but here refer to it as VAE-Sim.

Thus, the purpose of the present article is to describe our own implementation of a simple VAE and its use in molecular similarity measurements as applied, in particular, to the set of drugs, metabolites and natural products that we have been using previously [25; 91-96] as our benchmark for similarity metrics.

## Methods

We considered and tested grammar-based and junction-tree methods such as those used by Kajino [35], that exploited some of the ideas developed by Dai [97], Kusner [98] and by Jin and their colleagues [34]. However, our preferred method as described here used one-hot encoding as set out by Gómez-Bombarelli and colleagues [69]. We varied the number of molecules in the training process from ca 250,000 to over 6 million; the large number of possible hyperparameters would have led to a combinatorial explosion, so exhaustive search was (and is) not possible. The final architecture used here (shown in Fig 2C) required 6 days’ training on a 1-GPU machine. It involved a CNN encoder with the following layers (Fig 2C): **convolution (1D):** size (in-248=SMILES string length, 40 possible unique SMILES characters, out-9, kernel_size=9), **ReLU, convolution (1D)**: size (in-9, out-9, kernel_size=9) **ReLU, convolution (1D):** size (in-9, out-10, kernel_size=11) **ReLU, Linear (fully connected)**: size(140, latent_dims=100) **SeLU, with VAE mean**-Linear (fully connected): size(140, latent_dims=100) and **variance**-Linear (fully connected): size(140, latent_dims=100). For the decoder we used a **Reparametarization** (combined mean and sigma together) such that the output will be the same as the latent dimension (100 in our case), **Linear (fully connected):** size(latent_dims=100, latent_dims=100) **SeLU, RNN-GRU** (gated neural unit): size (hidden size=488, num_layers=3), **Linear (fully connected):** size(in-488=hidden_gru_size, out-248=SMILES length) **Softmax**. For the loss we used binary cross-entropy + KL-divergence. Neither dropout nor pooling were used. The optimiser was ADAM [99], the fixed learning rate 0.0001, parameters were initialised using the ‘Xavier uniform’ scheme [100], and a batch size of 128. This was implemented in Python using the Pytorch library (https://pytorch.org/). Most of the pre- and post-processing cheminformatics workflows were written in the KNIME environment (see [101]).

## Results

Autoencoders that use SMILES as inputs can return three kinds of outputs: (i) the correct SMILES output mirroring the input and/or translating into the input molecular structure (referred to as ‘perfect’), (ii) an incorrect output of a molecule different from the input but that is still legal SMILES (hence will return a valid molecule), referred to as ‘good’, and (iii) a molecule that is simply not legal SMILES. In practice, our VAE after training returned more than 95% valid SMILES in the test (holdout) set, so those that were invalid could simply be filtered out without significant loss of performance. Following training, each molecule (SMILES) could be associated with a normalised vector of 100 dimensions, and the Euclidean distance between them could be calculated.

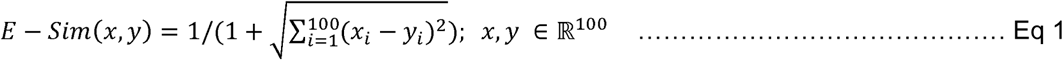

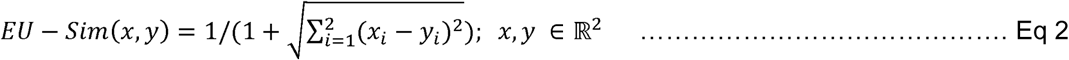

As previously [25], we compared the similarities between all drugs and all metabolites using the datasets made available in [91]. We here focus on just the MACCS and Patterned encodings of RDKit, and compare them with the normalised Euclidean distances according to the latent vector obtained from the VAE. As before, we rank ordered each drug in terms of its closest similarity to any metabolite. First, Fig 3A (reading from right to left) shows the Tanimoto similarities for the Patterned and MACCS fingerprints, as well as the VAE-Sim values as judged by two metrics. The first, labelled E-Sim (Eq 1), is the Euclidean similarity, based on the raw 100-dimensional hidden vectors, while the second, EU-Sim (Eq 2), used the Uniform Manifold Approximation and Projection (UMAP) dimension reduction algorithm [102; 103] based on the first two UMAP dimensions was used for purposes of visualisation; clearly, as with other encodings, they do not at all follow the same course, and one that may be modified according to the similarity measure used. Figures 3B and 3C show the ‘all-vs-all’ heatmaps for two of the encodings, indicating again that the VAE-Sim encoding falls away considerably more quickly, i.e. that similarities are judged in a certain sense more ‘locally’.

**Figure 3.**
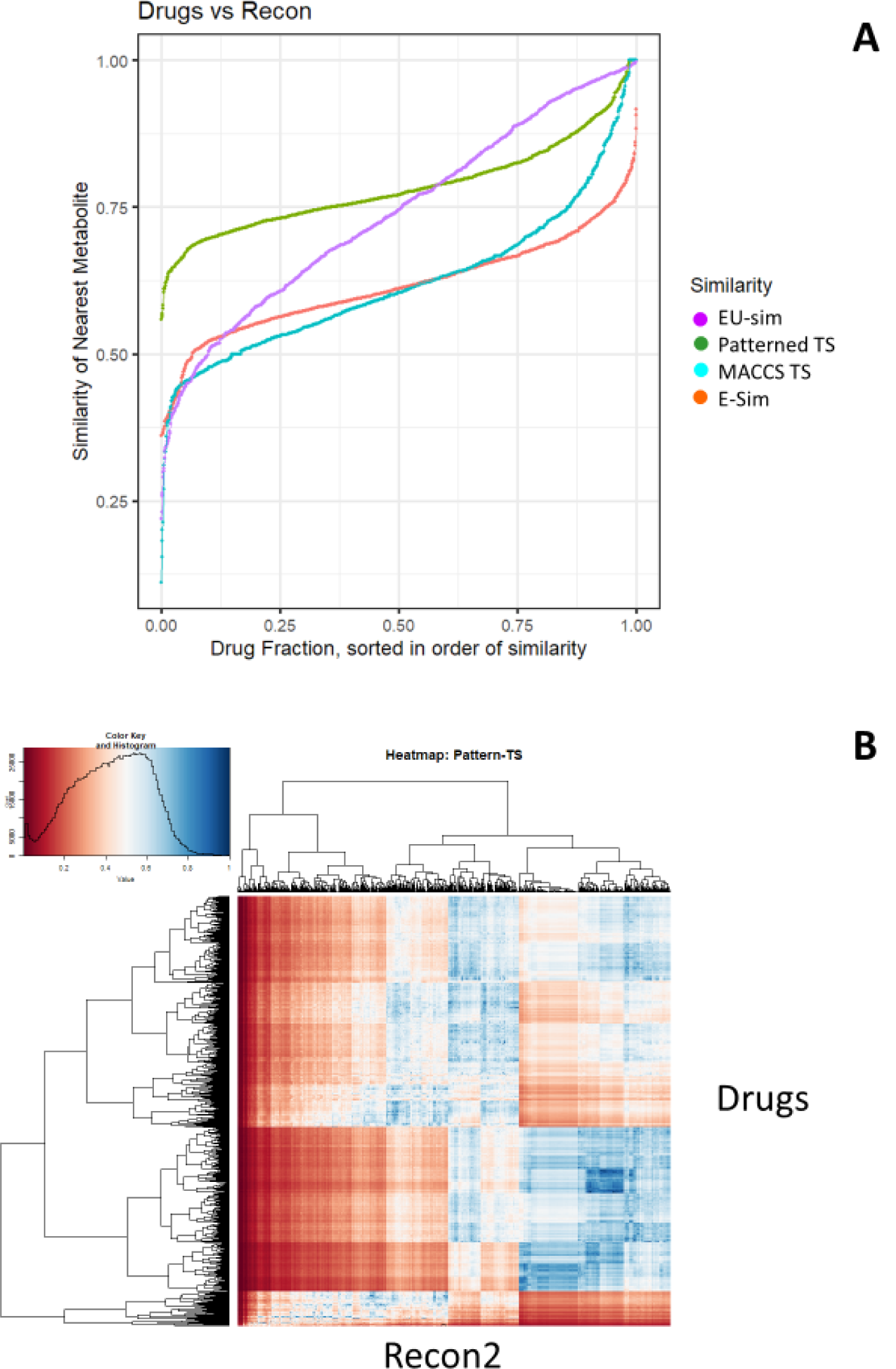
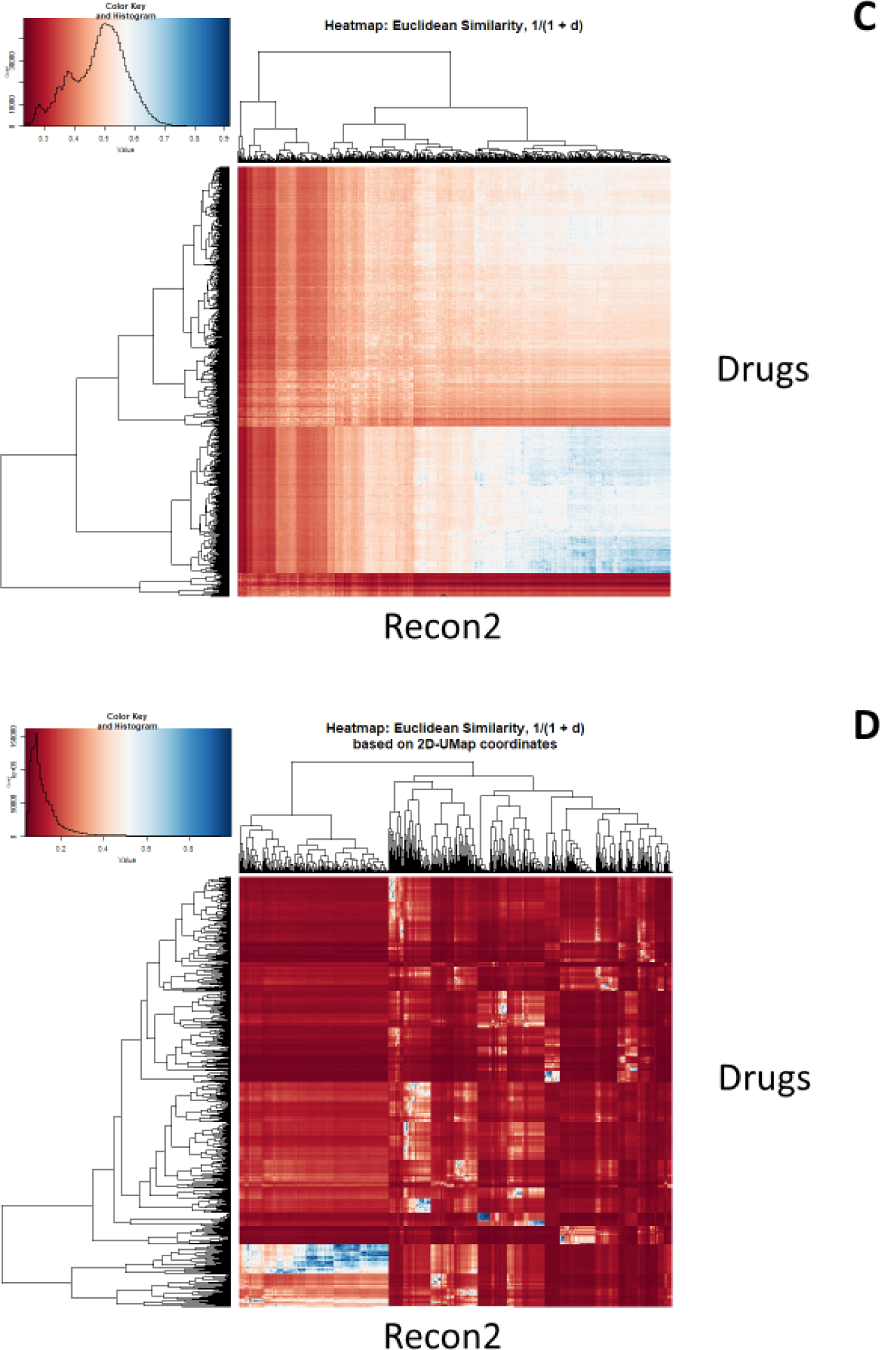
Top similarities between drugs and metabolites as judged by a fingerprint encoding (RDKit patterned) and our new VAE-Sim metric. **A**. Rank ordering. **B**. Heatmap for Tanimoto similarities using RDKit patterned encoding. **C**. Heatmap of Euclidean similarities E-Sim (Eq 1) for VAE-Sim in the 100-dimensional latent vector). **D** Heatmap of Euclidean similarities EU-Sim (Eq 2) for VAE-Sim in 2-dimensional UMAP space.

Figure 4A shows the Patterned similarity for the ‘most similar’ metabolite for each drug (using TS) compared to that for VAE-Sim (using Euclidean distance), while Figure 4B shows the same for the MACCS encoding. These again illustrate how the new encoding provides a quite different readout from the standard fingerprint encodings.

**Figure 4.**
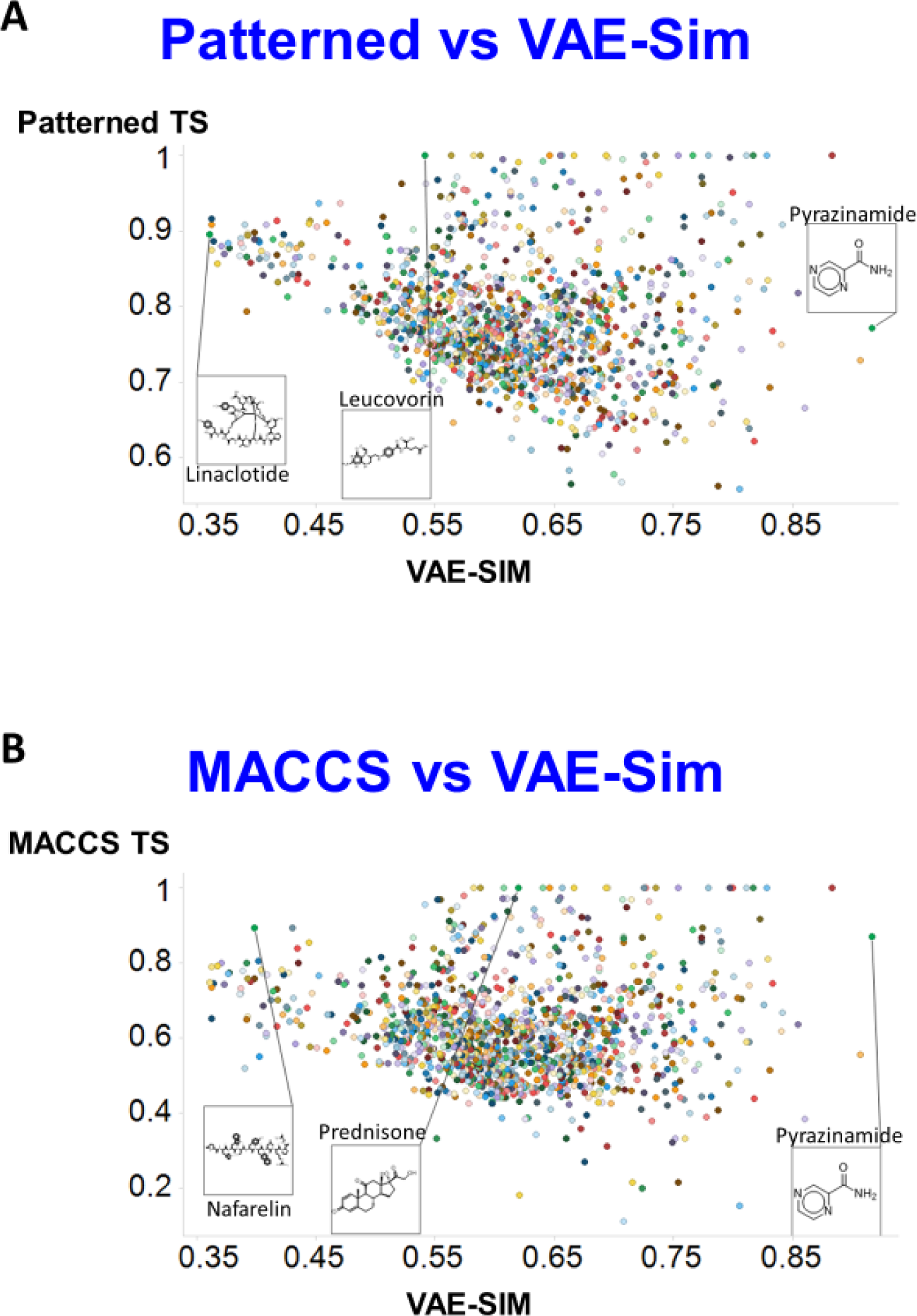
Comparison of similarities between two RDKit fingerprint methods and VAE-Sim Using Tanimoto similarity for fingerprints and Euclidean d_100_ similarity for VAE-Sim. **A**. Patterned encoding. **B**. MACCS encoding.

Finally, we used our new metrics to determine the similarity to clozapine of other drugs. Figure 5 shows the two similarity scores based on VAE-Sim, calculated as in Figure 3. Gratifyingly, and while structural similarity is, in part, in the eye of the beholder, a variety of structurally and functionally related antipsychotic drugs such as loxapine, mirtazapine and quetiapine were indeed among the most similar to clozapine, while others not previous considered (such as the antihistamines ketotifen and alcaftadine and the anti-inflammatory COX inhibitor ketorolac) were also suggested as being similar, providing support for the orthogonal utility of the new VAE-Sim metric. However, the rather promiscuous nature of clozapine binding (e.g. [104; 105]), and that of many of the other drugs (e.g. [106-112]), mean that this is not the place to delve deeper.

**Figure 5.**
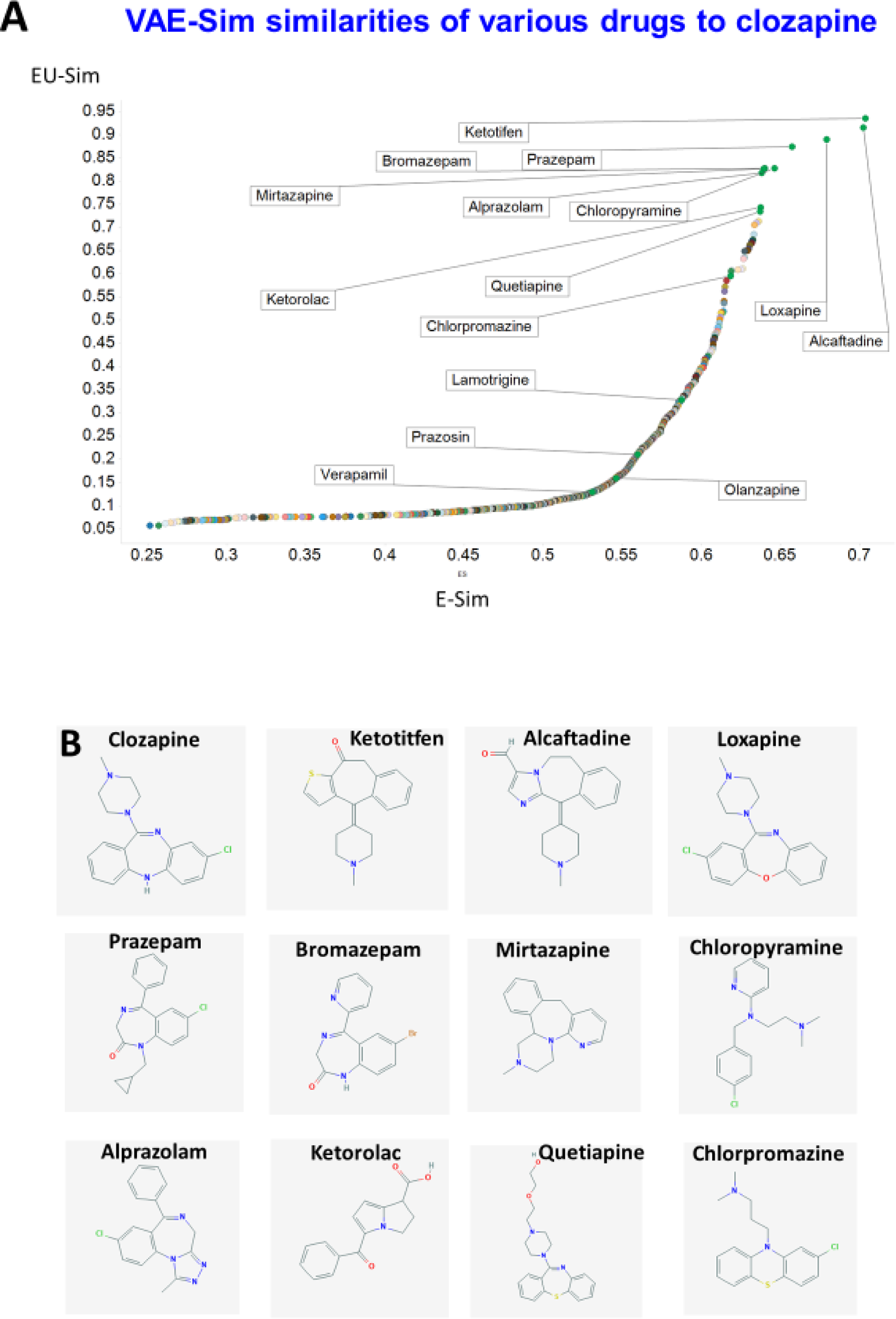
Similarity of drugs to clozapine as judged by the VAE. **A**. Rank order of Euclidean similarity in 100 dimensions (E-Sim) vs 2 UMAP dimensions (EU-Sim) as in Figure 3. Some of the ‘most similar’ drugs are labelled, as are some of those in Table 1. **B**. Structures of some of the drugs mentioned.

## Discussion

Molecular similarity is at the core of much of cheminformatics (e.g. [3; 8; 113-116]), but is an elusive concept. Our chief interest typically lies in supervised methods such as QSARs, where we use knowledge of paired structures and activities to form a model that allows us to select new structures with potentially desirable activities. Modern modelling methods such as feedforward artificial neural networks based on multilayer perceptrons are very powerful (and they can in fact fit any nonlinear function – the principle of “universal approximation” [117; 118]). Under these circumstances it is usually possible to learn a QSAR using any of the standard fingerprints. However, what we are focused on here is a purely unsupervised representation of the structures themselves (cf [37] which used substructures), and the question of which of these are the ‘most similar’ to a query molecule of interest. Such unsupervised methods may be taken to include any kinds of unsupervised clustering too (e.g. [119-123]). As with any kind of system of this type, the ‘closeness’ is a function of the weighting of any individual features, and it is perhaps not surprising that the different fingerprint methods give vastly different similarities, even when judged by rank order (e.g. [25] and above). One similarity measure that is independent of any fingerprint encoding is represented by the maximum common substructure (MCSS). However, by definition, the MCSS uses only part of a molecule; it is also computationally demanding [93; 94], such that ‘all-against-all’ comparisons such as those presented here are out of the question for large numbers of molecules.

Here we have leveraged a new method that uses only the canonical SMILES encoding of the molecules themselves, leading to its representation as a 100-element vector. Simple Euclidean distances can be used to obtain a metric of similarity that unlike MCSS is rapidly calculated for any new molecule, even against the entire set of molecules used in the development of the latent space.

In addition, unlike any of the other methods described, methods such as VAEs are <u>generative</u>: moving around in the latent space and applying the vector so created to the decoder allows for the generation of entirely new molecules (e.g. [41-45; 48; 50; 58; 60; 68; 69; 124]). This opens up a considerable area of chemical exploration, even in the absence of any knowledge of bioactivities.

### What determines the extent to which VAEs can generate novel examples?

The ability of variational autoencoders to generalise is considered to be based on learning a certain ‘neighbourhood’ around each of the training examples [72; 125], seen as a manifold of lower dimensionality than the dimensionality of the input space [51]. Put another way, “the reconstruction obtained from an optimal decoder of a VAE is a convex combination of examples in the training data” [126]. On this basis, an effect of training set size on the improvement of generalisation (here defined simply as being able to return an accurate answer from a molecule not in the training set) is to be expected, and our ability to generalise (as judged by test set error) improved as the number of molecules increased up to a few million. However, although we did not explore this, it is possible that our default architecture was simply too large for the smaller number of molecules, as excessive ‘capacity’ can cause a loss of generalisation ability [126]. This of course leaves open the details of the size and ‘closeness’ of that neighbourhood, how it varies with the encoding used (our original problem) and what features are used in practice to determine that neighbourhood. The network described here took nearly a week to train on a well-equipped GPU-based machine, and exhaustive analysis of hyperparameters was not possible. Consequently, because an understanding of the importance of local density will vary as a function of the position and nature of the relevant chemical space, we are not going to pursue them here. What is important is (i) that we could indeed learn to navigate these chemical spaces, and (ii) that the VAE approach admits a straightforward and novel estimation of molecular similarity.

## Acknowledgments

Present funding includes part of the EPSRC project SuSCoRD (EP/S004963/1), partly sponsored by AkzoNobel. DBK is also funded by the Novo Nordisk Foundation (grant NNF10CC1016517). Tim Roberts is funded by the NIHR UCLH Biomedical Research Centre.

